# Poly-*N*-acetylglucosamine mediates *Cutibacterium acnes* biofilm formation and biocide resistance

**DOI:** 10.1101/2023.10.10.558046

**Authors:** Jeffrey B. Kaplan

## Abstract

Biofilm formation likely plays an important role in the pathogenesis of implant-related infections caused by *Cutibacterium acnes*. Biofilms protect bacteria from antimicrobials and host defenses which makes biofilm-related infections difficult to treat. Here we demonstrate that the exopolysaccharide poly-*N*-acetylglucosamine (PNAG) contributes to *C. acnes* biofilm formation *in vitro*. By treating *C. acnes* cells and biofilms with the PNAG-degrading enzyme dispersin B, we found that PNAG mediates the attachment of *C. acnes* cells to polystyrene rods and the formation *C. acnes* biofilms in glass and polypropylene tubes. We further show that PNAG protects *C. acnes* biofilm cells from killing by tetracycline and benzoyl peroxide. PNAG may play an important role in biofilm formation, antibiotic tolerance, and virulence in this opportunistic pathogen.

## INTRODUCTION

*Cutibacterium acnes* is a Gram-positive bacterium that colonizes human skin. Although considered a beneficial commensal, *C. acnes* can sometimes cause invasive infections of the skin, soft tissue, cardiovascular system, and implanted medical devices (Achermann *et al*., 2014). *C. acnes* also contributes the pathogenesis of the common inflammatory dermatosis acne vulgaris (McLaughlin *et al*., 2019).

Poly-*N*-acetylglucosamine (PNAG) is an extracellular polysaccharide that mediates biofilm formation, antimicrobial resistance, host colonization, immune evasion, and stress tolerance in a wide range of Gramnegative and Gram-positive bacterial pathogens, as well as most fungi (Cywes-Bentley *et al*., 2013; Soliman *et al*., 2018). PNAG is an essential virulence factor for *Staphylococcus aureus* in mouse models of systemic infection (Kropec *et al*., 2005); for *Aggregatibacter actinomycetemcomitans* in a rat model of periodontitis (Shanmugam *et al*., 2015); for *Klebsiella pneumoniae* in a mouse model of intestinal colonization and systemic infection (Chen *et al*., 2014); and for *Actinobacillus pleuropneumoniae* during natural porcine pleuropneumonia infection in pigs (Subashchandrabose *et al*., 2013). Anti-PNAG antibodies have been shown to protect mice against local and/or systemic infections caused by *Streptococcus pyogenes, Streptococcus pneumoniae, Listeria monocytogenes, Neisseria meningitidis* serogroup B, and *Candida albicans* (Cywes-Bentley *et al*., 2013), suggesting that PNAG is an important virulence factor in these organisms.

Recent immunofluorescent microscopy studies using anti-PNAG antibodies showed that two *Cutibacterium* spp. strains, one of which was *C. acnes*, produced PNAG (Colette Cywes-Bentley and Gerald Pier, personal communication). However, no studies have investigated the function of PNAG in *C. acnes*. In this study we used the PNAG-degrading enzyme dispersin B to demonstrate that PNAG mediates the attachment of *C. acnes* cells to polystyrene rods as well as biofilm formation by *C. acnes* in glass and polypropylene tubes. We further show that PNAG protects *C. acnes* biofilm cells from killing by the antiacne agents tetracycline and benzoyl peroxide.

## MATERIALS AND METHODS

### Antimicrobial agents and enzymes

Tetracycline hydrochloride (Sigma-Aldrich; Catalog No. T7660) was dissolved at 10 mg/ml in distilled water, filter sterilized, and diluted in broth to the indicated concentra-tions. Benzoyl peroxide (TCI Chemicals; Catalog No. B3152) was dissolved at 10 mg/ml in dimethyl sulfoxide and diluted directly in broth. Sodium dodecyl sulfate (SDS; Catalog No. 428018) was purchased from Merck. Dispersin B was obtained from Kane Biotech (Winnipeg MB, Canada). Deoxyribonuclease I (Catalog No. DN25) was from Sigma-Aldrich.

### Bacterial strains and growth conditions

The *C. acnes* strains used in this study are listed in Table 1. All strains were isolated from patients with severe acne. Strains were maintained and enumerated on Tryptic Soy agar (BD). For broth cultures, bacteria were inoculated into filter-sterilized Tryptic Soy broth (BD) at ca. 10^6^ CFU/ml and incubated statically in an anaerobic atmosphere at 37°C for 72 h. Anaerobic conditions were created using a BD GasPak EZ Anaerobe sachet system. Bacteria were cultured in a 1-ml volume in 13 × 100 mm glass tubes or 15-ml conical-bottom polypropylene centrifuge tubes.

**TABLE 1.** Bacterial strains.

| Strain | Characteristics | Source* |
| --- | --- | --- |
| *C. acnes* HL072PA1 | Ribotype RT5 | BEI Resources |
| *C. acnes* HL043PA1 | Ribotype RT5 | BEI Resources |
| *C. acnes* HL086PA1 | Ribotype RT8, Erm<sup>r</sup> | BEI Resources |
| *C. acnes* HL036PA1 | Ribotype RT532 | BEI Resources |

\**C. acnes* strains were obtained through BEI Resources as part of the NIAID NIH Human Microbiome Project.

### Crystal violet binding assay

Biofilms cultured in glass tubes were rinsed vigorously with tap water and stained for 1 min with 1 ml of Gram’s crystal violet. Tubes were then rinsed with tap water to remove the unbound dye, air-dried, and photographed. To quantitate crystal violet binding, stained tubes were filled with 1 ml of 33% acetic acid, incubated at room temperature for 30 min, and mixed by vortex agitation. A volume of 200 μl of the dissolved dye was transferred to the well of a 96-well microtiter plate and its absorbance at 595 nm was measured in a microplate reader. Tubes containing sterile broth were incubated and processed along with the inoculated tubes to serve as controls. Biofilm inhibition was calculated using the formula (*A*595_Dispersin B_/*A*595_No enzyme_) × 100.

### Surface attachment assay

A suspension of *C. acnes* cells scraped from an agar plate (ca. 10^6^ CFU/ml in PBS) was aliquotted into three 15-ml centrifuge tubes (2.5 ml/tube). The first tube was left untreated to serve as a control. The second tube was supplemented with 20 μg/ml of dispersin B. The third tube was supplemented with 20 μg/ml of heat-inactivated dispersin B (95°C, 10 min). After 30 min at 37°C, the tubes were mixed by vortex agitation, and four 0.5-ml aliquots of each cell suspension were transferred to four separate 1.5-ml microcentrifuge tubes (0.5 ml/tube). A sterile 25-mm long polystyrene rod (1.5 mm diam; Plastruct Inc., Des Plaines IL, USA) was placed in each microcentrifuge tube. After 30 min, the rods were removed, rinsed with PBS to remove loosely adherent cells, and transferred to 15-ml conical centrifuge tubes containing 1 ml of PBS. Cells were detached from the rods by sonication, diluted, and plated on agar for CFU enumeration.

### Autoaggregation assay

*C. acnes* cells were scraped from an agar plate into 2 ml of PBS using a cell scraper. Aliquots of the cell suspension were treated with 50 μg/ml dispersin B or 100 μg/ml DNase I for 15 min. One aliquot of cells was left untreated as a control. A total of 300 μ□l of each cell suspension was transferred to a 0.5-ml polypropylene tube (model 6530; Corning), and the tube was then mixed by high-speed vortex agitation for 10 s, incubated statically for 15 min, and photographed.

### Treatment of biofilms with enzymes and detergent

Mature 72-h-old biofilms grown in glass tubes were rinsed vigorously with water and then treated with 1 ml of 100 μg/ml dispersin B, 100 μg/ml DNase I, or 1% SDS. After 15 min, tubes were rinsed vigorously with water and stained with crystal violet as described above.

### Benzoyl peroxide killing assay

Mature 72-h-old biofilms grown in glass tubes were treated directly with 20 μg/ml dispersin B for 15 min followed by 70 or 140 μg/ml benzoyl peroxide for 10 min. Biofilms were then detached from the tubes by sonication, diluted, and plated on agar for CFU enumeration.

### Tetracycline killing assay

Biofilms were cultured in glass tubes in broth supplemented with 20 μg/ml dispersin B and/or 0.2 μg/ml tetracycline (MIC = 0.5 μg/ml). After 72 h, biofilms were detached from the tubes by sonication, diluted, and plated on agar for CFU enumeration.

### Statistics and reproducibility of results

All experiments were performed in triplicate or quadruplicate tubes. All experiments were performed on at least two occasions with similar results. The significance of differences between means was calculated using a Student’s *t*-test. A *P-*value < 0.01 was considered significant.

## RESULTS

### Dispersin B inhibits *C. acnes* biofilm formation

The ability of four *C. acnes* strains to form biofilms in glass culture tubes was investigated using a crystal violet binding assay (Fig. 1). Two of the four strains tested (HL043PA1 and HL036PA1) formed strong biofilms as evidenced by the high amount of bound crystal violet dye.

**Figure 1.**
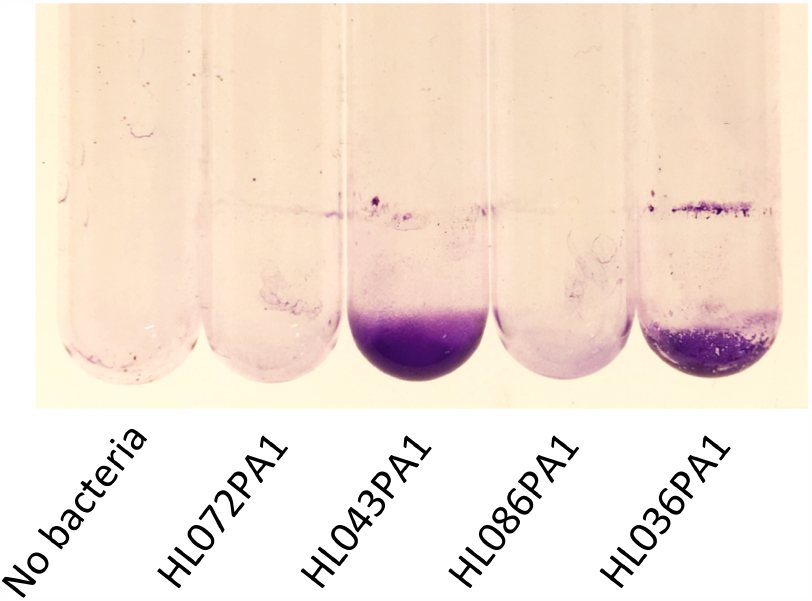
***C. acnes* biofilm formation in glass tubes. Cultures were incubated for 3 d, rinsed vigorously with water, stained with crystal violet for 1 min, re-rinsed with water, dried, and photographed. Strain names are indicated below**.

To determine whether dispersin B inhibits *C. acnes* biofilm formation, biofilm-forming strains HL043PA1 and HL036PA1 were incubated in unsupplemented broth or in broth supplemented with 100 μg/ml dispersin B (Fig. 2). Dispersin B significantly inhibited biofilm formation by both strains as evidenced by the lower amount of bound crystal violet dye in tubes supplemented with the enzyme. Quantitation of bound dye for strain HL034PA1 yielded a biofilm inhibi-tion value of 99% (*P* < 0.001; Fig. 3). These results suggest that PNAG contributes to *C. acnes* biofilm formation.

**Figure 2.**
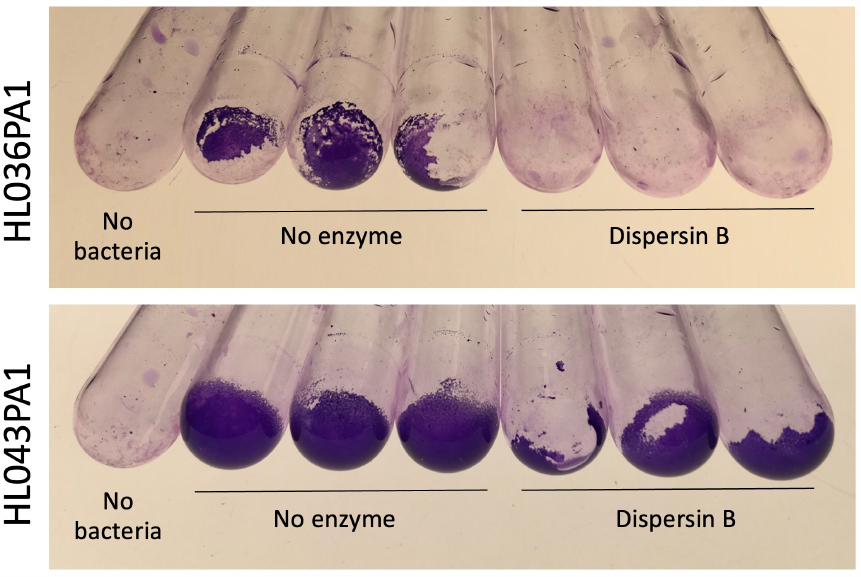
***C. acnes* biofilm formation in the absence or presence of 100 μg/ml dispersin B. Triplicate tubes for each condition and one control tube are shown. Strain names are indicated at the left**.

**Figure 3.**
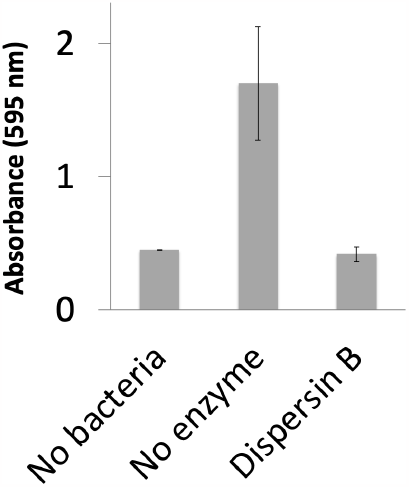
**Quantitation of *C. acnes* HL036PA1 biofilm formation in the absence or presence of 100 μg/ml dispersin B. Values show mean and range for triplicate tubes for each condition**.

### Dispersin B inhibits attachment of *C. acnes* cells to surfaces

Photographs of HL043PA1 and HL036PA1 glass culture tubes taken directly after incubation (prior to rinsing and crystal violet staining) revealed the presence of thin biofilms along the sides of the tubes that were absent in tubes supplemented with dispersin B (Fig. 4). This phenomenon was also evident for HL043PA1 and HL036PA1 cultured in conical-bottom polypropylene centrifuge tubes (Fig. 5 and data not shown). In both types of tubes, however, a significant amount of cell clumping was observed at the bottom of the tube, even in the presence of the enzyme. These observations suggest that dispersin B may inhibit the attachment of *C. acnes* biofilms and biofilm cells to surfaces rather than inhibiting biofilm formation itself. To test this hypothesis, untreated and dispersin B-treated *C. acnes* planktonic cells were incubated in the presence of polystyrene rods and the number of cells that attached to the rods after 30 min was enumerated (Fig. 6). Significantly fewer dispersin B-treated cells attached to the rods than untreated cells, whereas cells treated with heat-inactivated dispersin B attached to the rods at the same level as untreated cells. These results suggest that PNAG contributes to *C. acnes* surface attachment.

**Figure 4.**
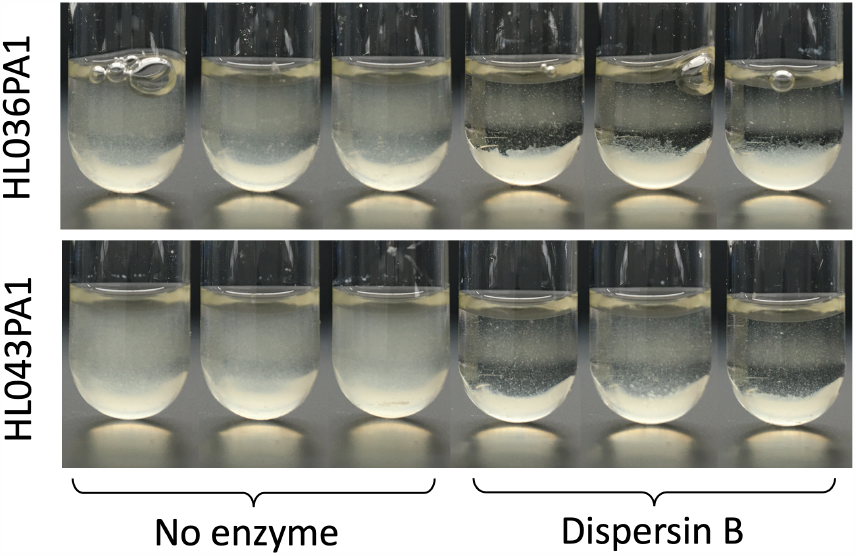
***C. acnes* growth in the absence or presence of 100 μg/ml dispersin B. Triplicate tubes for each condition are shown. Strain names are indicated at the left**.

**Figure 5.**
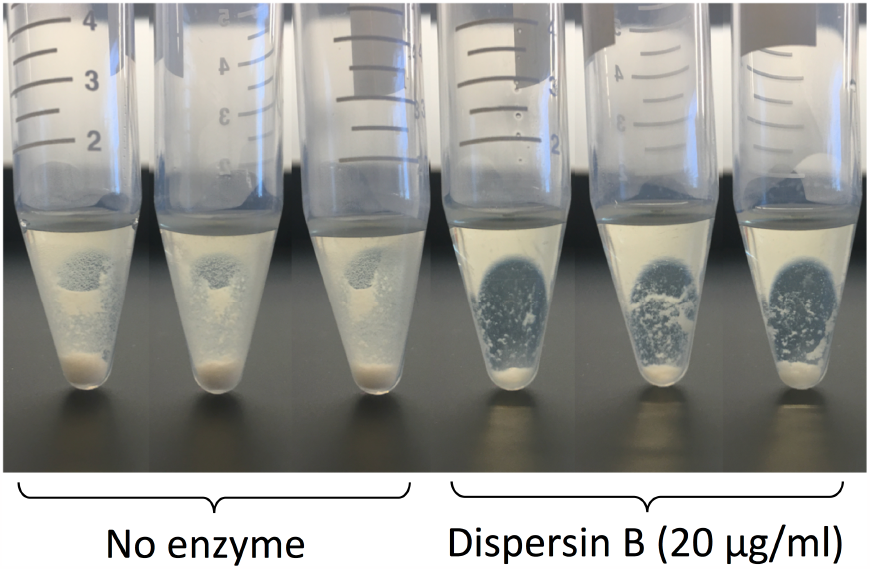
***C. acnes* HL036PA1 growth in polypropylene centrifuge tubes in the absence or presence of 100 μg/ml dispersin B. Triplicate tubes for each condition are shown**.

**Figure 6.**
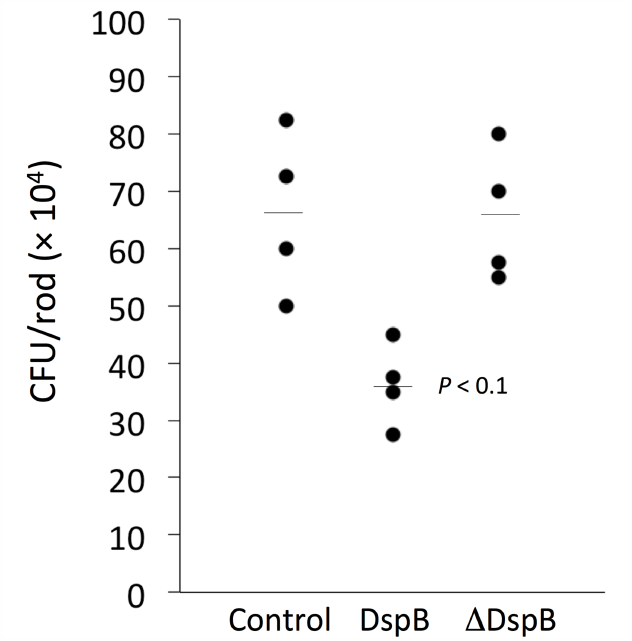
**Attachment of *C. acnes* HL036PA1 cells to polystyrene rods. Some cells were treated with dispersin B (DspB) or heat-inac1vated dispersin B (ΔDspB) prior to contac1ng the rods. Each dot represents one rod. Horizontal lines indicate means**.

### DNase I, but not dispersin B, inhibits *C. acnes* autoaggregation

Autoaggregation (also termed intercellular adhesion) often plays a role in biofilm formation. To test whether PNAG contributes to *C. acnes* autoaggregation, *C. acnes* HL036PA1 cells were treated with dispersin B or DNase I for 30 min, mixed by vortex agitation, transferred to a microcentrifuge tube, allowed to settle for 15 min, then photographed (Fig. 7). Untreated control cells and dispersin B-treated cells settled to the bottom of the tube, whereas DNase I-treated cells remained in suspension. These findings suggest that extracellular DNA, but not PNAG, contributes to *C. acnes* autoaggregation.

**Figure 7.**
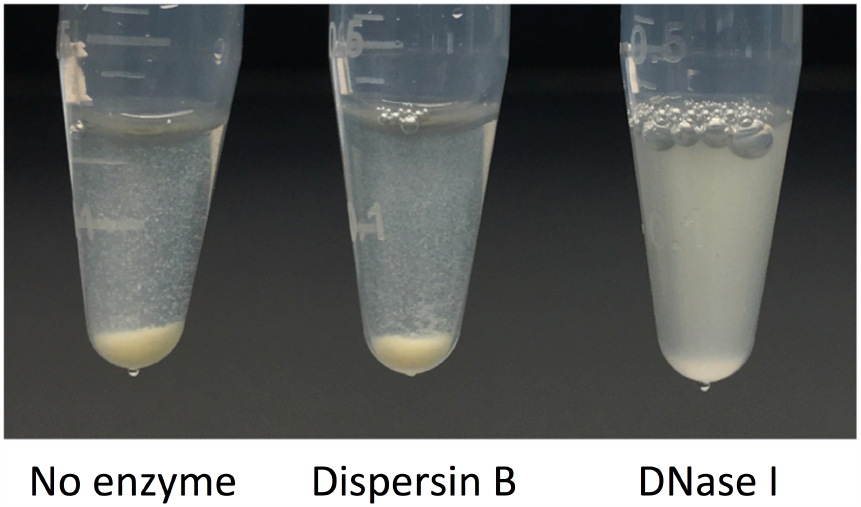
**Autoaggregation of untreated, dispersin B-treated, and DNase I-treated *C. acnes* HL036PA1 cells**.

### Detachment of *C. acnes* biofilms by enzymes and detergents

To further investigate the composition of the *C. acnes* biofilm matrix, mature 72-h-old HL043PA1 biofilms were treated with dispersin B, DNase I, or 1% SDS for 15 min, and then stained with crystal violet to visualize the biofilm remaining after treatment (Fig. 8). DNase I and SDS efficiently detached the mature biofilms, suggesting that extracellular DNA and proteinaceous adhesins may contribute to biofilm stability in mature *C. acnes* biofilms.

**Figure 8.**
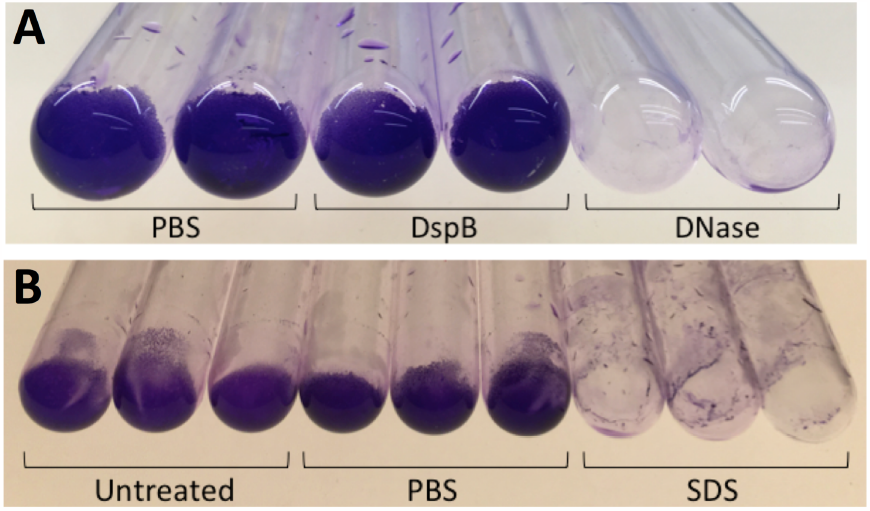
**Detachment of *C. acnes* HL043PA1 biofilms by enzymes and detergents. Panel A shows biofilms treated with phosphate buffered saline (PBS)**, **dispersin B (DspB) or DNase I (DNase) prior to crystal violet staining. Duplicate tubes for each condition are shown. Panel B shows untreated biofilms and biofilms treated with PBS or 1% SDS prior to crystal violet staining. Triplicate tubes for each condition are shown**.

### Dispersin B sensitizes *C. acnes* biofilms to benzoyl peroxide killing

Mature 72-h-old HL043PA1 biofilms were treated directly with 20 μg/ml dispersin B for 15 min followed by 70 or 140 μg/ml benzoyl peroxide for 10 min. The biofilms were then detached from the tubes by sonication, diluted, and plated on agar for CFU enumeration. Control experiments showed that dispersin B alone did not kill *C. acnes* cells or inhibit their growth (data not shown). Treatment of *C. acnes* biofilms with 70 or 140 μg/ml benzoyl peroxide alone resulted in a 1-log reduction in *C. acnes* CFUs, while pre-treatment of biofilms with dispersin B increased benzoyl peroxide killing by approximately 0.5 log (*P* < 0.001; Figs. 9 & 10). These findings suggest that PNAG protects *C. acnes* biofilm cells from killing by benzoyl peroxide.

**Figure 9.**
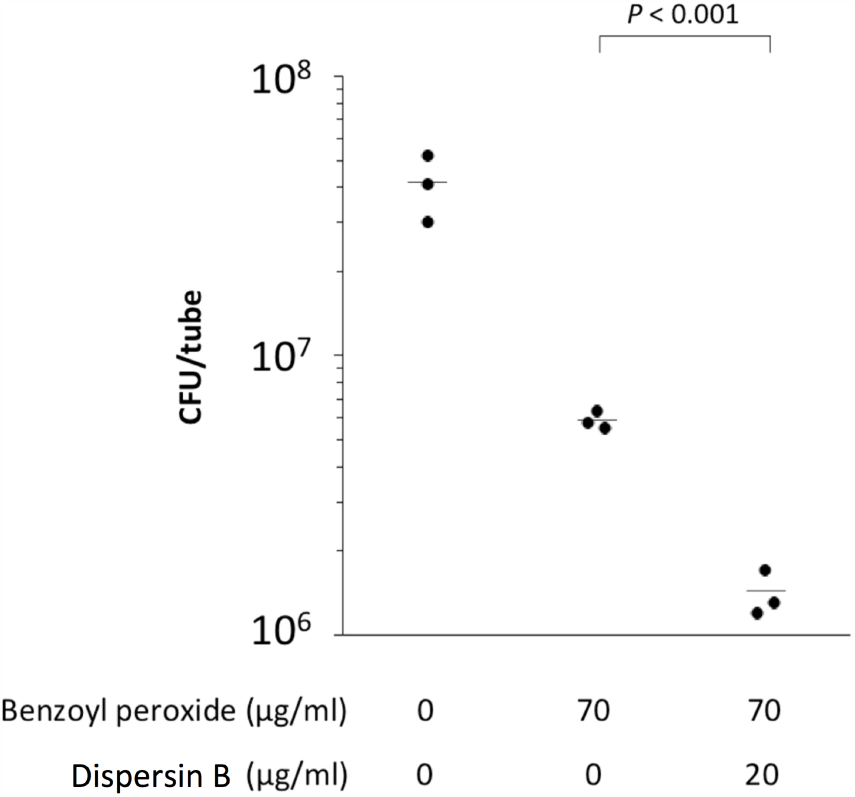
**Dispersin B sensi1zes *C. acnes* HL043PA1 biofilms to killing by benzoyl peroxide. Mature bofilms were treated with 0 or 20 μg/ml dispersin B for 15 min followed by 70 μg/ml benzoyl peroxide for 10 min. Bio-films were then detached from the tubes by sonication, diluted, and plated on agar for CFU enumertion. Each dot represents one tube**.

**Figure 10.**
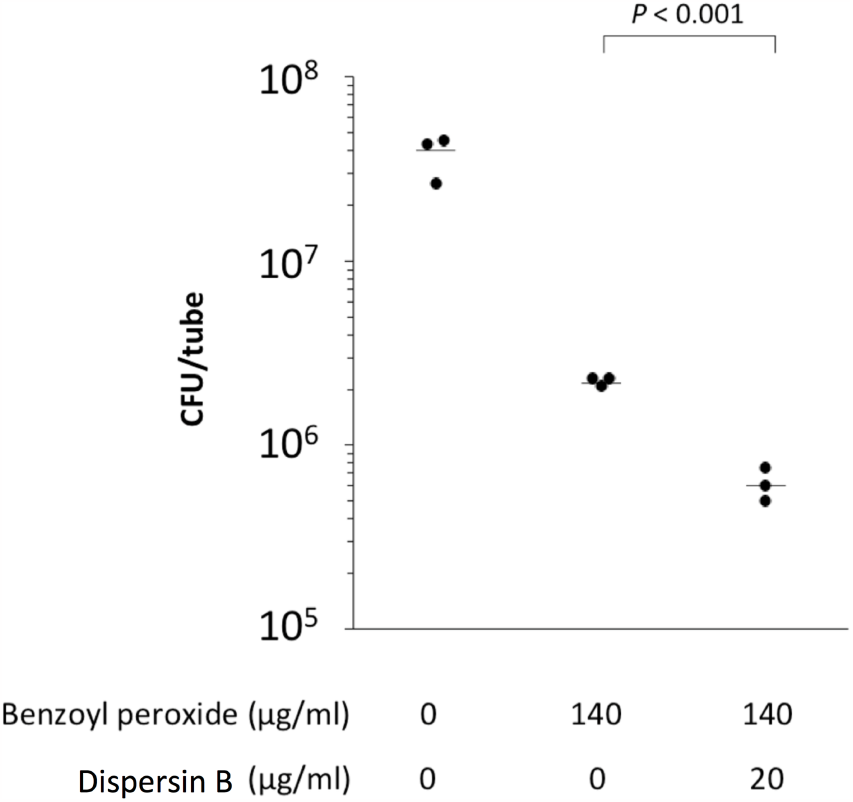
**Dispersin B sensi1zes *C. acnes* HL043PA1 bio-films to killing by benzoyl peroxide. Mature bofilms were treated with 0 or 20 μg/ml dispersin B for 15 min followed by 140 μg/ml benzoyl peroxide for 10 min. Biofilms were then detached from the tubes by sonication, diluted, and plated on agar for CFU enumertion. Each dot represents one tube.**

### Dispersin B decreases antibiotic tolerance in *C. acnes* biofilms

The effect of dispersin B on the tolerance of HL043PA1 biofilms to tetracycline was measured by culturing biofilms in broth supplemented with 20 μg/ml dispersin B and/or 0.2 μg/ml tetracycline (MIC = 0.5 μg/ml). Significantly fewer *C. acnes* cells were recovered from tubes supplemented with dispersin B plus tetracycline compared to tubes supplemented with tetracycline alone (Fig. 11). These findings suggest that PNAG may contribute to antibiotic tolerance *C. acnes* biofilms.

**Figure 11.**
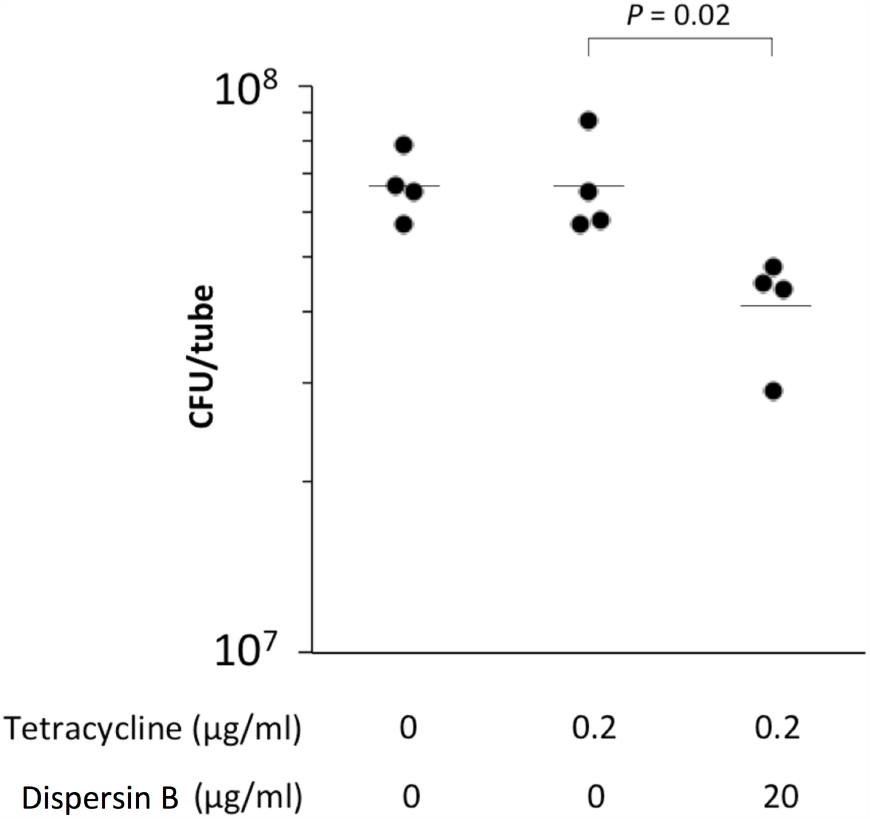
**Dispersin B sensi1zes *C. acnes* HL043PA1 biofilms to killing by tetracycline. Biofilms were cultured in the presence of 20 μg/ml dispersin B and/or 0.2 μg/ml tetracycline (MIC = 0.5 μg/ml). After 72 h, biofilms were detached from the tubes by sonication, diluted, and plated on agar for CFU enumertion. Each dot represents one tube**.

## DISCUSSION

Biofilms are densely-packed layers of bacterial cells growing attached to a tissue or surface (Costerton *et al*., 1999). Bacteria in a biofilm are encased in a sticky, self-synthesized polymeric matrix that holds the cells together in a mass, attaches them to the underlying surface, and protects them from killing by antimicrobial agents and host immunity. *C. acnes* biofilms have been observed on implanted medical devices and in acne lesions *in vivo* (Bayston *et al*., 2006; Jahns *et al*., 2012). *In vitro* studies on the adhesive components of the *C. acnes* biofilm matrix revealed the presence of polysaccharides, proteinaceous adhesions, and extra-cellular DNA (Coeyne *et al*., 2022). In the present study we found that *C. acnes* biofilms were detached by SDS and DNase I, consistent with the presence of proteinaceous adhesins and extracellular DNA in the biofilm matrix. One previous study (Okuda *et al*., 2018) showed that dispersin B did not inhibit biofilm formation by five strains of *C. acnes* isolated from cardiac pacemakers when cultured in polystyrene microtiter plate wells, suggesting that PNAG is not an adhesive component of *C. acnes* biofilms. In the present study, however, we found that dispersin B inhibited biofilm formation by two *C. acnes* strains isolated from acne patients when cultured in glass tubes, confirming that PNAG contributes to *C. acnes* biofilm cohesion in some strains. The discrepancy between Okuda *et al*. (2018) and the present study may be due to strain differences or to the fact that dispersin B exhibits different anti-biofilm activities depending on the shape and size of the culture vessel (Izano *et al*., 2007b).

Dispersin B is the only known enzyme that hydrolyzes the β-1-6 glycosidic linkages of PNAG (Cywes-Bentley *et al*., 2013). Because of its high specificity, dispersin B can be used to detect the presence of PNAG (Eddenden & Nitz, 2022). In the present study we found that purified dispersin B not only inhibited *C. acnes* biofilm formation, but also inhibited attachment of *C. acnes* cells to polystyrene rods and sensitized *C. acnes* biofilms to killing by benzoyl peroxide and tetracycline, suggesting that PNAG plays a role in these processes. These antibiofilm activities are consistent with those exhibited by dispersin B against other species of bacteria (Ganeshnarayan *et al*., 2009; Itoh *et al*., 2005; Izano *et al*., 2007a; Parise *et al*., 2007). Because of its antibiofilm activity against *C. acnes*, it is possible that dispersin B may be a useful agent for presurgical skin antisepsis during shoulder arthroplasty surgery or for the treatment and prevention of acne vulgaris.

Our results demonstrate that PNAG contributes to *C. acnes* surface attachment, biofilm formation, and biocide resistance *in vitro*, suggesting that PNAG may be an important *C. acnes* virulence factor. PNAG may enable *C. acnes* to form biofilms and colonize epithelial surfaces and hair follicles, and to resist killing by antimicrobial agents and host immunity. PNAG may also function as a biological glue that holds corneocytes together to form microcomedones (Burkhart & Burkhart, 2007).

## ACKNOWLEDGEMENTS

The author thanks Colette Cywes-Bentley and Gerald Pier (Harvard Medical School) for providing data on PNAG production in *Cutibacterium* spp.; Kane Biotech Inc. (Winnipeg, Manitoba, Canada) for providing dispersin B; BEI Resources (Manassas, Virginia, USA) for providing bacterial strains; and Katie DeCicco-Skinner (American University) for continued support.

